# Altitude-mediated niche partitioning between *Dacus bivittatus* and *Dacus punctatifrons* along an elevational transect in the Uluguru Mountains, Tanzania

**DOI:** 10.64898/2026.04.21.720022

**Authors:** Maulid W. Mwatawala, Joseph O. Ruboha, Jackline Bakengesa, Mwajuma K. Zinga, Marc De Meyer

## Abstract

Understanding how fruit fly species partition resources along environmental gradients is important for predicting pest pressure under changing climatic conditions. The population ecology of *Dacus bivittatus* (Bigot) and *Dacus punctatifrons* (Karsch) (Diptera: Tephritidae) was examined across six sites spanning 526–1,650 m above sea level in the Uluguru Mountains, Tanzania, over eight years (2004–2012). A total of 2,200 weekly trap records were aggregated into 292 site-month observations and standardised as flies per trap per day (FTD). *Dacus bivittatus* showed strong seasonal structuring (H = 43.03, p < 0.001), with abundance peaking during the cool dry season (June–August), whereas *D. punctatifrons* showed no clear seasonal pattern. Both species declined significantly with increasing altitude (ρ = −0.308 and −0.769, respectively; p < 0.001), but the decline was steeper for *D. punctatifrons*. Species dominance shifted along the gradient: *D. punctatifrons* dominated warm lowland conditions (>24 °C), whereas *D. bivittatus* prevailed at elevations above approximately 569 m. Seasonal niche overlap declined markedly with altitude, indicating increasing temporal segregation between the species in cooler environments. These findings demonstrate that altitude structures ecological divergence between two closely related fruit fly pests and provide a basis for site-specific monitoring and climate-sensitive pest forecasting in tropical mountain agroecosystems.

## Introduction

Fruit flies (Diptera: Tephritidae) are among the most important pests of horticultural crops worldwide because larval feeding within fruits reduces yield and market quality and may trigger quarantine and trade restrictions (1–3). In tropical and subtropical regions, their impact is severe because warm conditions permit prolonged breeding and rapid population growth (4). In sub-Saharan Africa, cucurbit crops such as cucumber (*Cucumis sativus* L.), watermelon (*Citrullus lanatus* [Thunb.] Matsum. & Nakai), pumpkin, and squash (*Cucurbita* spp.) are important sources of food, nutrition, and income, particularly for smallholder farmers (5,6). Recent agroecological studies in eastern-central Tanzania showed that, in the absence of effective control measures, infestations can become severe resulting in substantial production losses (5,7).

Despite their economic importance, seasonality and climate–population relationships are well understood for only a limited number of African fruit fly species studied over long periods and across multiple localities (e.g., 8–11). Among these, *Dacus bivittatus* (Bigot) and *Dacus punctatifrons* (Karsch) are widely distributed across the Afrotropical region and are frequently associated with cultivated and wild cucurbit hosts (4,12,13). Most previous information on the two species come from short-term trapping, fruit-incubation studies, or broad faunistic surveys involving multiple fruit fly taxa (14,15).

Relatively more ecological information is available for *D. punctatifrons* than other cucurbit-infesting *Dacus* species. Seasonal patterns have been reported for *D. punctatifrons* (16), while *D. bivittatus* remains mainly documented through general surveys (15,17,18).

Fruit fly populations are regulated by interacting climatic and ecological drivers, particularly temperature, rainfall, humidity, and host plant availability, all of which affect development, survival, and reproductive performance (4,19,20). Long-term, standardised trapping data are essential for accurately describing seasonality, interannual fluctuations, and climate–population relationships in indigenous *Dacus* species.

Environmental gradients such as altitude may further structure these dynamics by altering temperature, rainfall, humidity, vegetation structure, and host-resource distribution across space. Along mountain transects, such gradients can produce marked shifts in insect abundance and species composition. At the same time, heterogeneous agro-landscapes may intensify these patterns by creating mosaics of cultivated habitats, semi-natural vegetation, and refuge areas. In the Uluguru Mountains and comparable tropical systems, fruit fly assemblages vary along elevational gradients, with climate and host availability jointly influencing occurrence and relative abundance (17,21).

Against this background, the present study used a long-term parapheromone-based trapping dataset to examine the population ecology of *D. bivittatus* and *D. punctatifrons* along an altitudinal transect in the Uluguru Mountains of Tanzania. By combining multi-year abundance records with climatic variables and agroecological zone classification, the study aimed to assess seasonal dynamics, inter-annual variability, climatic sensitivity, and species overlap across a lowland–highland gradient (22,23).

This study tested the hypothesis that environmental gradients along the Uluguru Mountains structure the ecological relationship between *D. bivittatus* and *D. punctatifrons*. Specifically, it was expected that the abundance of both species would decline with increasing altitude because cooler highland conditions generally constrain fruit fly activity and population growth, but that the rate of decline would differ between species because of differences in ecological tolerance. Based on previous studies (21,24), *D. punctatifrons* was expected to be more closely associated with warmer lowland environments, whereas *D. bivittatus* was expected to persist more successfully at higher elevations.

## Material and methods

### Agroecological zones and site distribution along an altitudinal gradient

Monitoring of *D. bivittatus* and *D. punctatifrons* was conducted in the Morogoro Region of Tanzania along a lowland–highland altitudinal transect spanning four agroecological zones, characterised by differences in altitude, land use, and host-plant composition along the eastern Uluguru Mountains (25–27). This zonation captures systematic changes in cropping systems and host availability along the altitudinal transect, comprising the lowland, mid-elevation transition, sub-montane, and montane zones.

The lowland zone is characterised by warm conditions and diverse agricultural mosaics dominated by tropical fruit systems. The mid-elevation transition zone comprises mixed smallholder production systems. The sub-montane zone is dominated by staple-based production systems with relatively limited availability of fruit hosts. The montane zone is defined by cooler conditions and agroforestry systems dominated by temperate fruit production (27).

### Sampling design and landscape-level trapping

Six monitoring sites spanning 500–1650 m a.s.l. were established along the transect to capture a continuous agroecological gradient from lowland tropical systems to montane agroforestry landscapes (Fig. 1). Across this transect, representatives of Cucurbitaceae and Solanaceae, including cultivated and wild species, were present. Cultivated cucurbits included cucumber (*Cucumis sativus* L.), melon (*Cucumis melo* L.), watermelon (*Citrullus lanatus* (Thunb.) Matsum. & Nakai), pumpkin (*Cucurbita moschata* (Duchesne) Duchesne ex Poir.), luffa (*Luffa acutangula* (L.) Roxb.), and bitter gourd (*Momordica charantia* L.), alongside wild hosts including *Momordica* cf. *trifoliata*, *M.* cf. *foetida*, *Cucumis dipsaceus*, and related species.

**Fig. 1.**
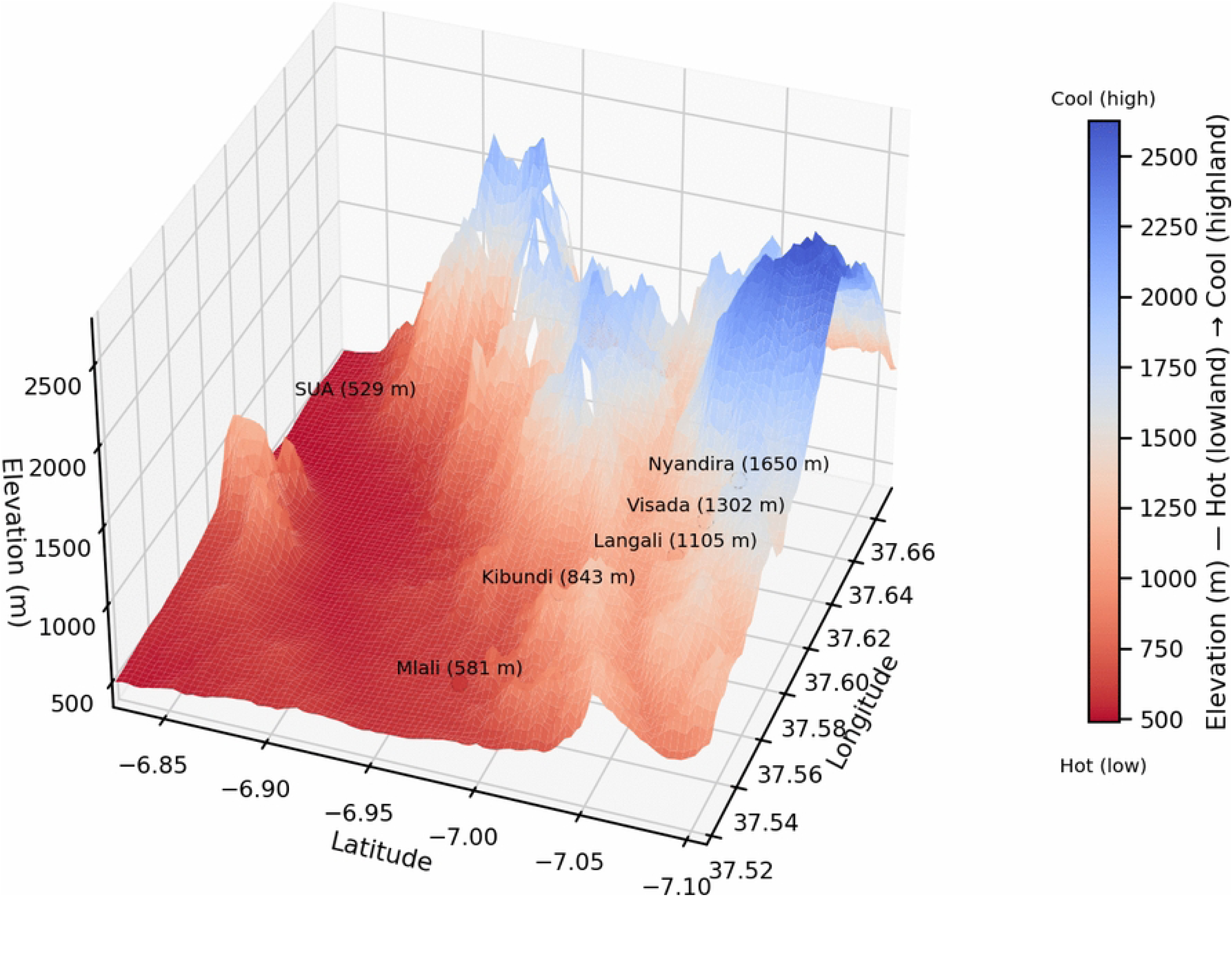
3D elevation surface around the six monitoring localities (SRTM DEM), highlighting the lowland–highland gradient.

Within this gradient, SUA (526 m a.s.l.) and Mlali (581 m a.s.l.) represent lowland agro-mosaics supporting diverse cropping systems alongside cucurbits. Kibundi (843 m a.s.l.) marks the mid-elevation transition, characterised by mixed smallholder systems combining tropical fruit trees *Mangifera indica* L., *Citrus* spp. L., and Annonaceae with crops in Poaceae and Fabaceae. Langali (1105 m a.s.l.) and Visada (1302 m a.s.l.) define the sub-montane zone, dominated by Poaceae and Fabaceae with limited fruit trees. Nyandira (1650 m a.s.l.) represents a cooler montane agroforestry system characterised by temperate fruit trees *Malus domestica* Borkh., *Prunus persica* (L.) Batsch, and *Prunus domestica* L. intercropped with *Coffea arabica* L. (see also 8,21,28).

### Host availability and crop phenology

In the study area, cucurbits are grown throughout both wet and dry seasons under continuous and staggered smallholder production systems, resulting in overlapping crop growth stages across the year. Fields at different phenological stages co-occur, and both cultivated and wild hosts are present within the landscape.

*D. bivittatus* and *D. punctatifrons* are largely stenophagous, being primarily associated with cucurbit hosts. Under these conditions, host availability is continuous (5), limiting the definition of discrete phenological windows. Consequently, crop phenology was not included as an explicit variable, and host availability is interpreted at the landscape level.

### Trap deployment and monitoring

Adult male *D. bivittatus* and *D. punctatifrons* were monitored using modified McPhail traps baited with cue-lure (Scentry Cie, Billings, MT, USA). Traps were suspended at approximately 2 m height in host trees. Each trap contained a single cue-lure plug and dichlorvos (DDVP; Scentry Cie) as a killing agent. The use of cue-lure, a potent male parapheromone, enables attraction of flies from considerable distances, allowing sampling of populations from the surrounding landscape. Trapping procedures followed protocols used by Mwatawala et al. (8) and Geurts et al. (21,28).

Monitoring at the SUA orchard began in mid-October 2004 with three traps placed in mango, citrus, and guava trees. In January 2005, two additional traps were added, one on mango and another on citrus, bringing the total to five traps. At Nyandira, monitoring also began in October 2004 with one trap placed in a peach tree, and a second trap was added in January 2005 in a plum tree. Monitoring at other sites along the altitudinal transect (Mlali, Kibundi, Langali, and Visada) commenced in October 2008, with three traps deployed at each site, mostly placed on mango trees.

Sampling was not continuous at all sites due to logistical constraints. At SUA, monitoring was continuous except for a break between November 2007 and March 2008. At Nyandira, there was a prolonged gap from October 2004 to September 2005, followed by several shorter interruptions. These periods were treated as months without sampling rather than months with zero captures.

Traps were inspected at regular intervals. During each inspection, traps were opened and all captured insects were removed. The number of *D. bivittatus* and *D. punctatifrons* captured per trap was recorded together with the number of days since the previous inspection, allowing abundance to be expressed as flies per trap per day (FTD).

At SUA, traps were inspected approximately once per week. At Nyandira and other transect sites, traps were deployed for one week every four weeks following a rotational schedule. After each sampling round, traps were removed and randomly reinstalled among host trees within the same site during the next cycle to minimise location bias.

Lure plugs and DDVP strips were replaced every four weeks. All captured specimens were transported to the laboratory at SUA for identification and data recording.

### Identification and taxonomic validation

Specimens were sorted from other insects and identified to species level by examining morphological characteristics under a stereomicroscope, using standard taxonomic guides (1,29,30).

### Climate data and gap management

Climatic data were obtained from several sources because data availability differed between the study sites. At SUA, monthly rainfall (mm), mean air temperature (°C), and mean relative humidity (%) were obtained from the Tanzania Meteorological Authority (TMA) station in the Morogoro area located about 1 km from SUA orchard. Temperature and relative humidity at the site were also recorded using iButton data loggers (Maxim Integrated, Sunnyvale, CA, USA).

At the other locations, air temperature in the orchards was recorded with iButton loggers and summarised as monthly averages. Relative humidity was not recorded with data loggers at these sites. Because reliable long-term rainfall data were not available, monthly rainfall values were obtained from spatial climate datasets for the sites.

When climate records were incomplete, additional data were obtained from reanalysis and gridded datasets. Monthly temperature and rainfall data were obtained from the ERA5 reanalysis using Google Earth Engine (31). WorldClim v2 bioclimatic variables were used to describe long-term climate conditions. These datasets were used only to fill missing data and support predictions and did not replace TMA or on-site measurements where these were available.

### Abundance index and temporal aggregation

The abundance of fruit flies was standardised as flies per trap per day (FTD) to account for variation in trapping effort. For each inspection interval *i*, FTD was calculated by dividing the total *D. bivittatus* and *D. punctatifrons* count (C*_i_*) by the number of traps in use (T*_i_*) multiplied by the number of days since the previous inspection (D*_i_*):

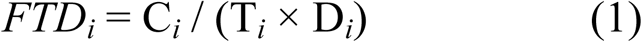

Monthly mean FTD for calendar month *m* was then computed as a trap-day weighted average: the sum of flies assigned to month *m* divided by the total trap-days occurring in that month. If an inspection interval crossed a month boundary, captures were apportioned between months in proportion to days spent in each month (equivalently assuming a constant daily capture rate within the interval). Monthly uncertainty was reported as the standard error of the inspection-interval FTD values that contributed trap-days to that month.

Summing daily FTD across a month produces a cumulative monthly catch per trap; in this study, monthly mean FTD is reported to facilitate comparisons across months and sites.

### Statistical analyses

All analyses were conducted in Python using standard statistical libraries (32–36). Seasonal, annual, and altitudinal differences in FTD were first assessed using non-parametric tests (Kruskal–Wallis) with pairwise Mann–Whitney comparisons where appropriate. Relationships between FTD and continuous environmental variables (altitude, temperature, rainfall, and relative humidity) were evaluated using Spearman’s rank correlations.

To identify independent effects of environmental predictors, multiple regression models were fitted for each species using log-transformed FTD values [log (FTD + 1)]. Finally, Tweedie generalised linear models (GLMs) were used to evaluate the combined influence of weather variables and agroecological zone on FTD while accommodating the right-skewed and zero-inflated distribution of trap catches. A preliminary analysis showed a strong negative correlation between altitude and temperature (Spearman’s ρ = −0.79). To avoid collinearity, agroecological zones were treated as a categorical landscape variable while retaining ecological interpretability (37).

### Species divergence and niche overlap

Species dominance patterns were examined using a log₂ dominance ratio calculated for each site-month record:

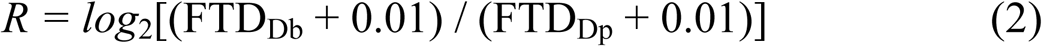

where positive values indicate *D. bivittatus* dominance and negative values indicate *D. punctatifrons* dominance. Niche overlap between the two species was quantified using Schoener’s D and Pianka’s O along three axes: altitudinal distribution, seasonal distribution, and seasonal patterns within sites. Values range from 0 (no overlap) to 1 (complete overlap). Overlap was also evaluated annually to assess temporal changes in niche differentiation.

## Results

### General temporal and spatial patterns of abundance

We observed clear spatial variation in trap catches of *D. bivittatus* and *D. punctatifrons* along the altitudinal transect (Fig. 2). Abundance was highest in the lowland zone, particularly at SUA, and declined toward the transition, sub-montane, and montane zones. In these higher zones, both species were present at low densities, with FTD rarely exceeding 0.6. We found that altitude significantly affected abundance of both species (Abundance declined with elevation, more steeply for *D. punctatifrons* than for *D. bivittatus* (Fig. 2)). Site-level comparisons supported this pattern. At the lowland site SUA (526 m), *D. punctatifrons* was more abundant than *D. bivittatus*. Above 843 m, within the transition and sub-montane zones, the pattern reversed, with *D. bivittatus* dominating. Only at Mlali (581 m), also in the lowland zone, did the two species occur at similar abundance (Fig. 2).

**Fig. 2.**
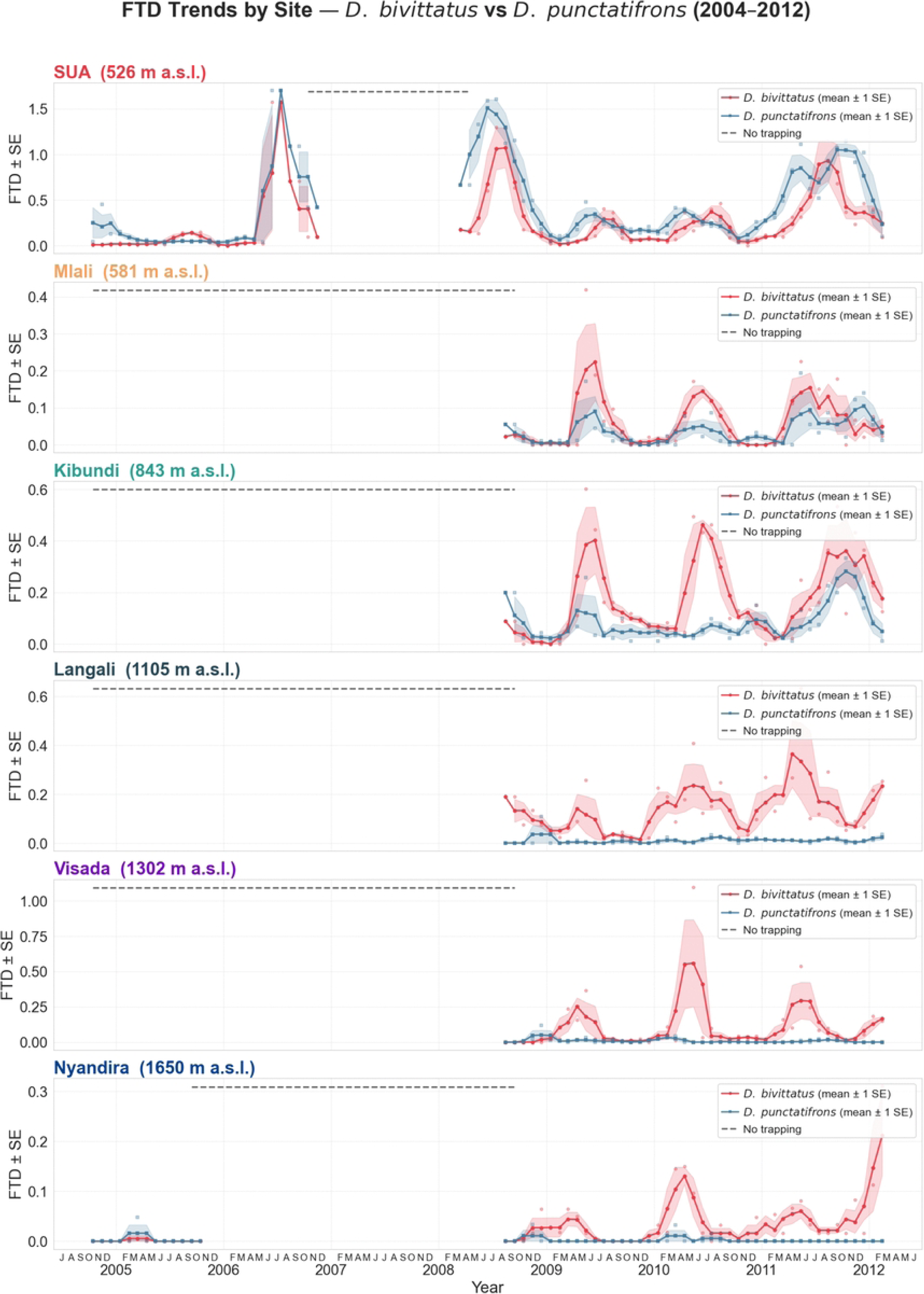
Spatial and temporal variation in trap catches of *D. bivittatus* and *D. punctatifrons* along the Uluguru Mountains transect.

### Temporal dynamics of abundance

#### Seasonal variation

We also observed clear differences in seasonal patterns between *D. bivittatus* and *D. punctatifrons* when data were pooled across sites and years (Fig. 3). *D. bivittatus* showed strong seasonal variation, with abundance peaking during the cool dry season (June–August). In contrast, *D. punctatifrons* showed no significant monthly variation, indicating a more uniform distribution throughout the year (Fig. 3). A more concentrated seasonal pattern was observed for *D. bivittatus* (Table 1), characterised by higher Markham concentration and a shorter activity period. In contrast, greater seasonal evenness and a longer activity window were observed for *D. punctatifrons*.

**Fig. 3.**
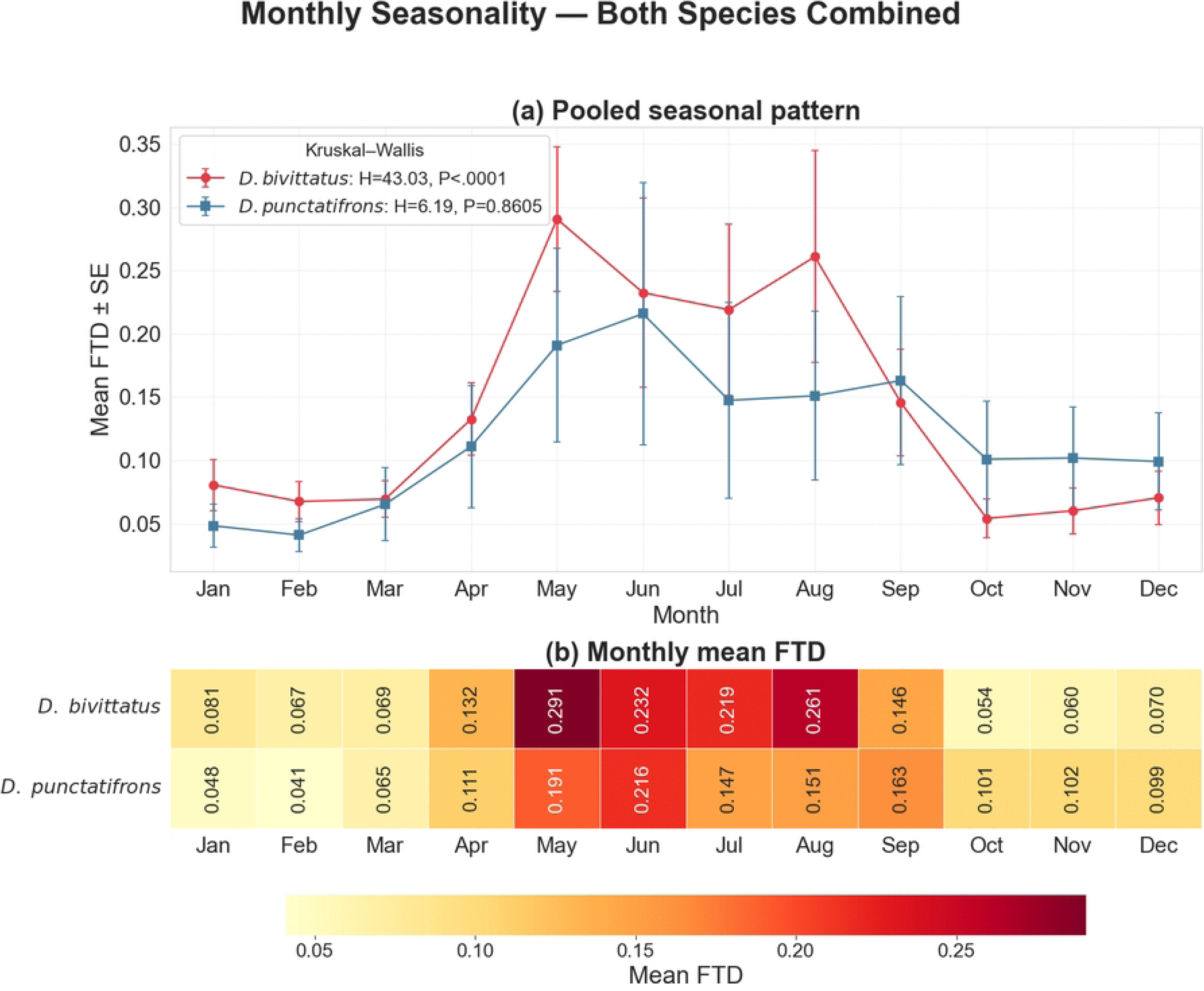
Seasonal abundance profiles of *D. bivittatus* and *D. punctatifrons* pooled across sites and years.

**Table 1.** Seasonal phenology of *D. bivittatus* and *D. punctatifrons* along the Uluguru Mountains transect.

| Species | Site | Peak month | Trough month | Active months | Seasonal concentration (%) | Evenness (J') |
| --- | --- | --- | --- | --- | --- | --- |
| *D. bivittatus* | SUA | Aug | Feb | 4 | 54.05 | 0.852 |
|  | Mlali | May | Oct | 2 | 60.91 | 0.791 |
|  | Kibundi | May | Mar | 4 | 35.71 | 0.910 |
|  | Langali | May | Oct | 1 | 26.95 | 0.919 |
|  | Visada | May | Oct | 1 | 69.07 | 0.722 |
|  | Nyandira | Feb | Oct | 4 | 51.40 | 0.866 |
|  | All sites | May | Oct | 5 | 37.50 | 0.930 |
| *D. punctatifrons* | SUA | Jun | Feb | 5 | 28.62 | 0.958 |
|  | Mlali | May | Mar | 1 | 30.22 | 0.877 |
|  | Kibundi | May | Feb | 7 | 22.19 | 0.954 |
|  | Langali | Dec | Oct | 2 | 16.51 | 0.857 |
|  | Visada | Dec | Jun | 1 | 24.43 | 0.855 |
|  | Nyandira | Mar | Jan | 1 | 48.63 | 0.352 |
|  | All sites | Jun | Feb | 6 | 27.11 | 0.959 |

*Note.* Active months indicate the number of months with $\geq 50\%$ of peak FTD. Seasonal concentration (%) represents the Markham concentration index describing the strength of seasonal aggregation, and J' represents Shannon evenness of monthly abundance.

#### Inter-annual variation

Year-to-year variation in abundance was evident for both species, although the magnitude of change differed between them. Abundance of *D. bivittatus* increased from the early monitoring years to a peak FTD around 2008, declined briefly in 2009, and then increased again toward the end of the study period (Fig. 4). A broadly similar annual pattern was observed for both species; however, *D. bivittatus* became more abundant than *D. punctatifrons* in the later years, particularly in 2010. A marginal year × species interaction was detected, indicating a gradual divergence between the two populations over time (Fig. 4).

**Fig. 4.**
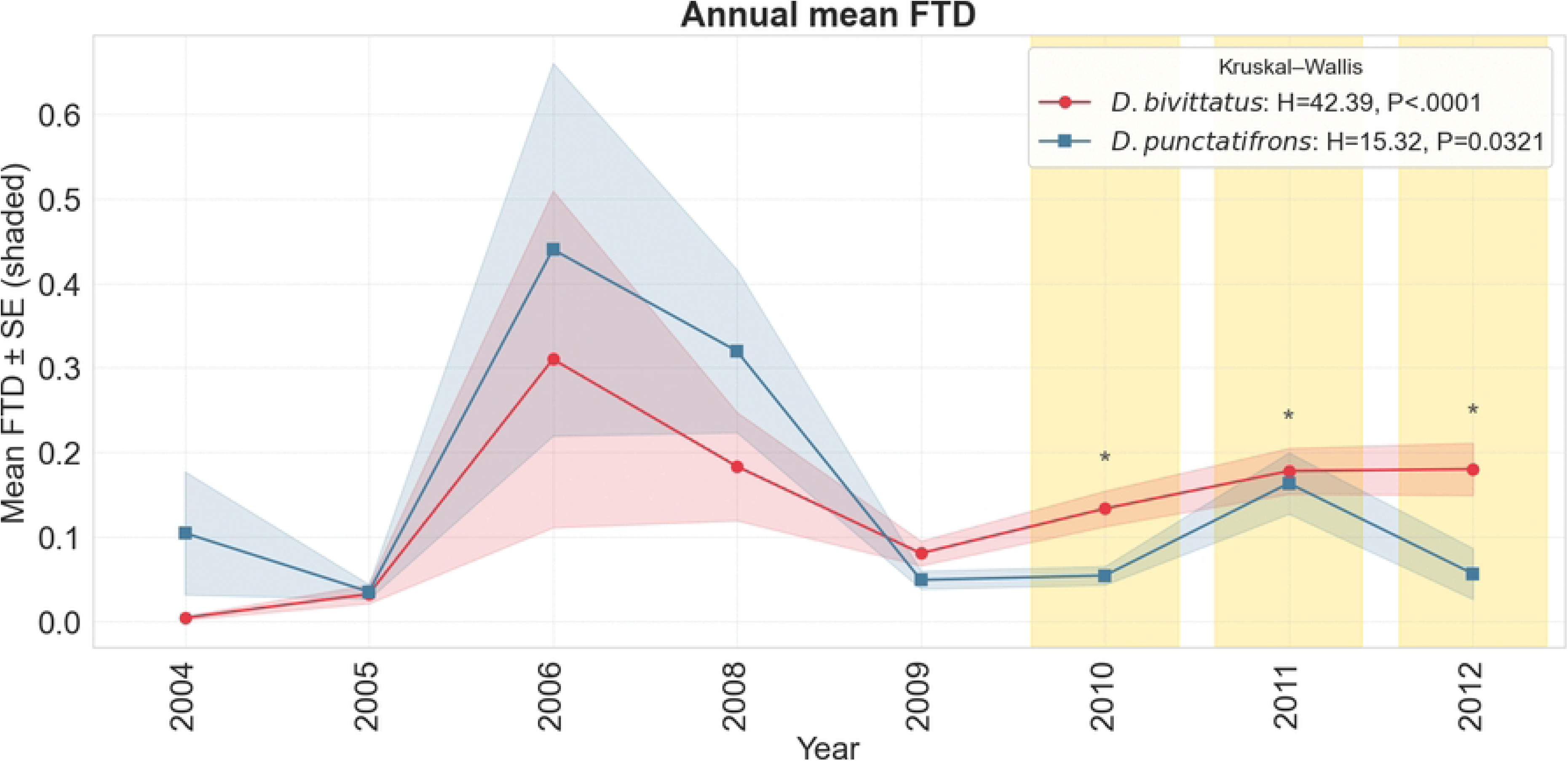
Inter-annual variation in mean FTD of *D. bivittatus* and *D. punctatifrons* from 2004–2012. Annual mean trap catches pooled across monitoring sites illustrate year-to-year population fluctuations and increasing divergence between the species in later years.

**Fig. 5.**
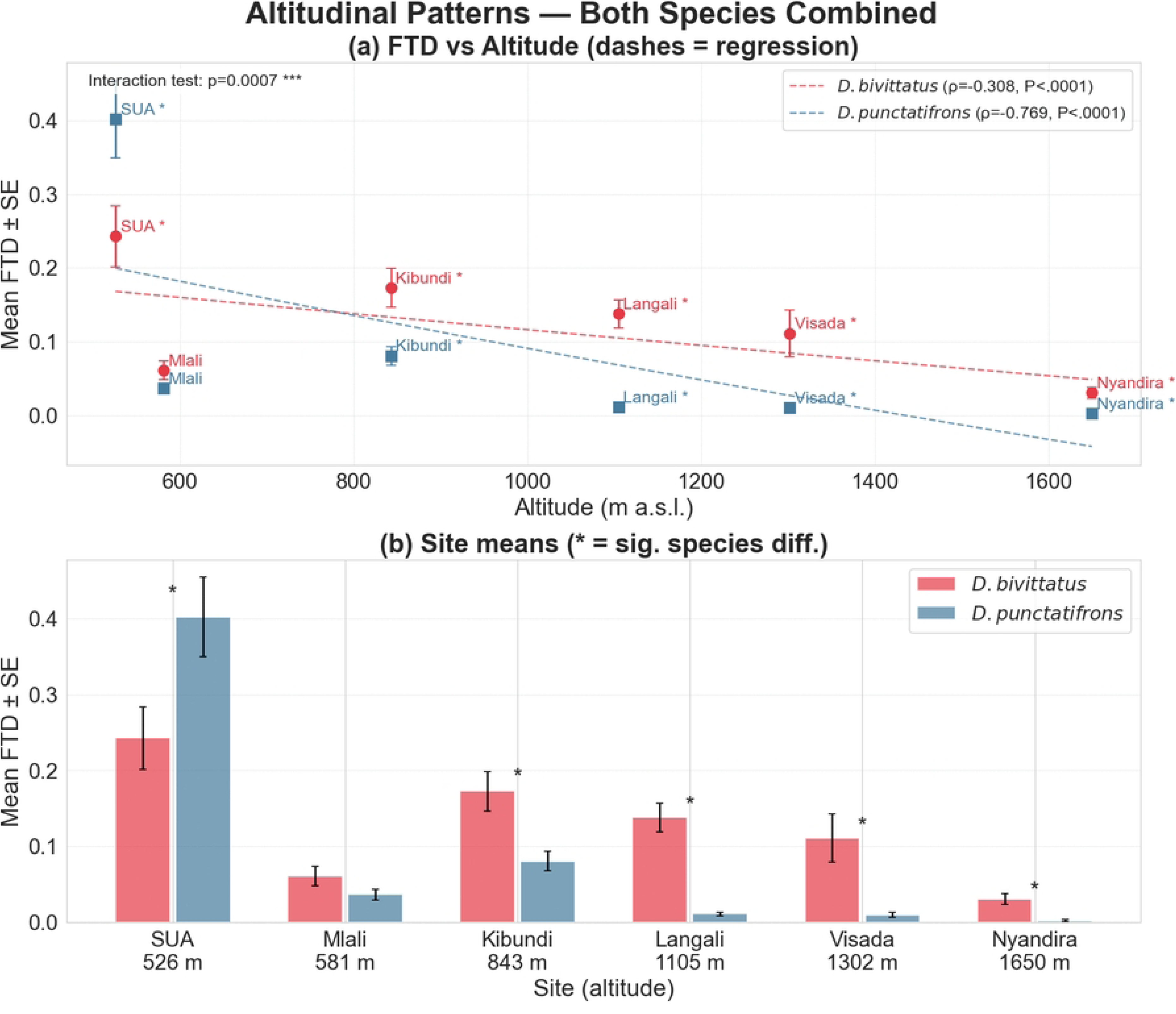
Altitudinal distribution of *D. bivittatus* and *D. punctatifrons* along the Uluguru Mountains transect. (a) Relationship between FTD and elevation; (b) mean site-level FTD of the two species.

### Environmental and landscape drivers of abundance

#### Altitudinal gradient

Across sites, weather variables showed contrasting relationships with the two species. *D. punctatifrons* abundance was positively associated with temperature and negatively associated with rainfall. In contrast, *D. bivittatus* showed no significant association with temperature and only a weak relationship with rainfall. Relative humidity was not significantly correlated with abundance for either species (Fig. 6). Multiple regression models yielded similar patterns, explaining more variance for *D. punctatifrons* than for *D. bivittatus*. Temperature was a significant predictor only for *D. punctatifrons*. A likelihood-ratio test confirmed that the two species responded differently to climatic variables.

**Fig. 6.**
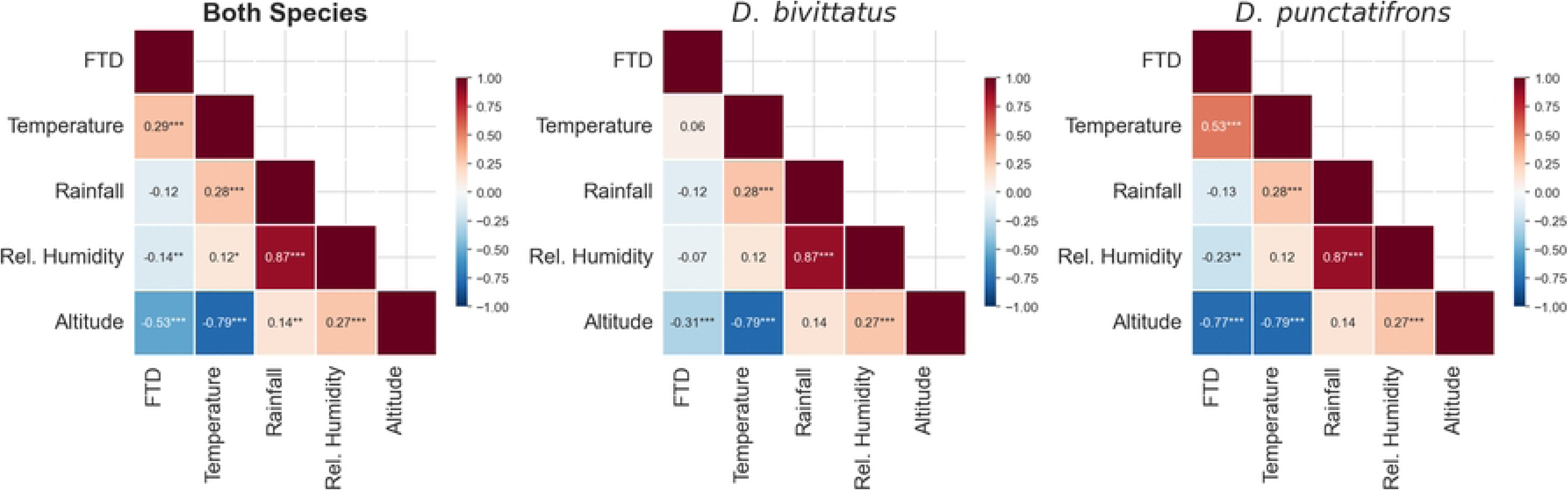
Relationships between weather variables and fruit fly abundance. Scatterplots showing associations between FTD and temperature, rainfall, and relative humidity for *D. bivittatus* and *D. punctatifrons*.

### Landscape effects on abundance

Generalised linear models showed strong effects of agroecological zones on abundance (Fig. 7). For *D. bivittatus*, abundance declined from the lowland zone to the montane zone, with reductions of 40% at mid-elevation, 74% in the sub-montane zone, and 97% in the montane zone. Temperature also had a significant effect within zones. For *D. punctatifrons*, the decline along the gradient was even stronger, with abundance reduced by 75% at mid-elevation, 97% in the sub-montane zone, and 99.6% in the montane zone. In contrast, weather variables contributed little once zonal differences were accounted for.

**Fig. 7.**
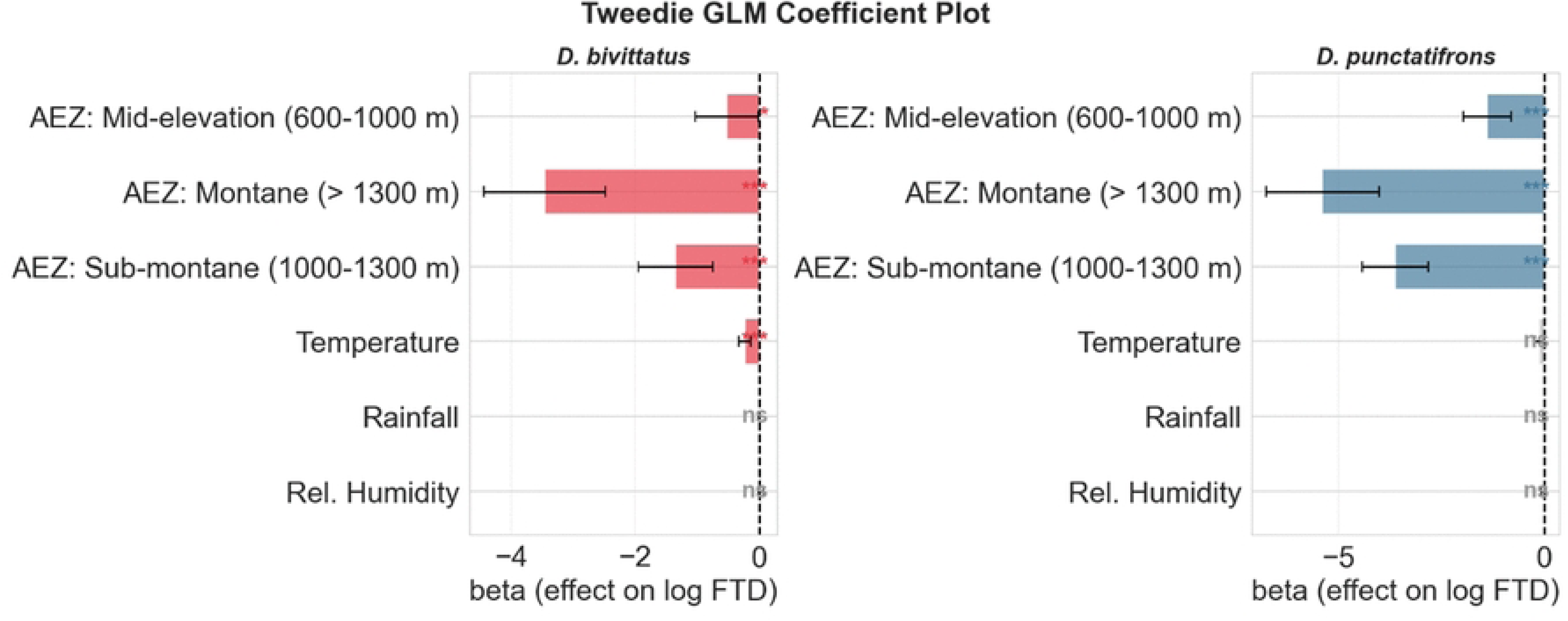
Effects of agroecological zones on fruit fly abundance from Tweedie GLMs. Coefficient estimates with 95% confidence intervals for predictors influencing FTD of *D. bivittatus* and *D. punctatifrons*.

### Species dominance and niche overlap

#### Dominance patterns along the gradient

A clear shift in species dominance occurred along the altitudinal gradient (Fig. 8). *D. punctatifrons* dominated in the lowland zone, particularly at SUA, whereas *D. bivittatus* became increasingly dominant through the transition and sub-montane zones, reaching its peak at Langali (≈ +3.0). Linear interpolation placed the dominance crossover at approximately 569 m above sea level, corresponding to the lower transition zone. A similar shift was observed along the temperature axis. LOWESS smoothing indicated a transition at approximately 24.1°C, above which *D. punctatifrons* dominated and below which *D. bivittatus* was more abundant (Fig. 8).

**Fig. 8.**
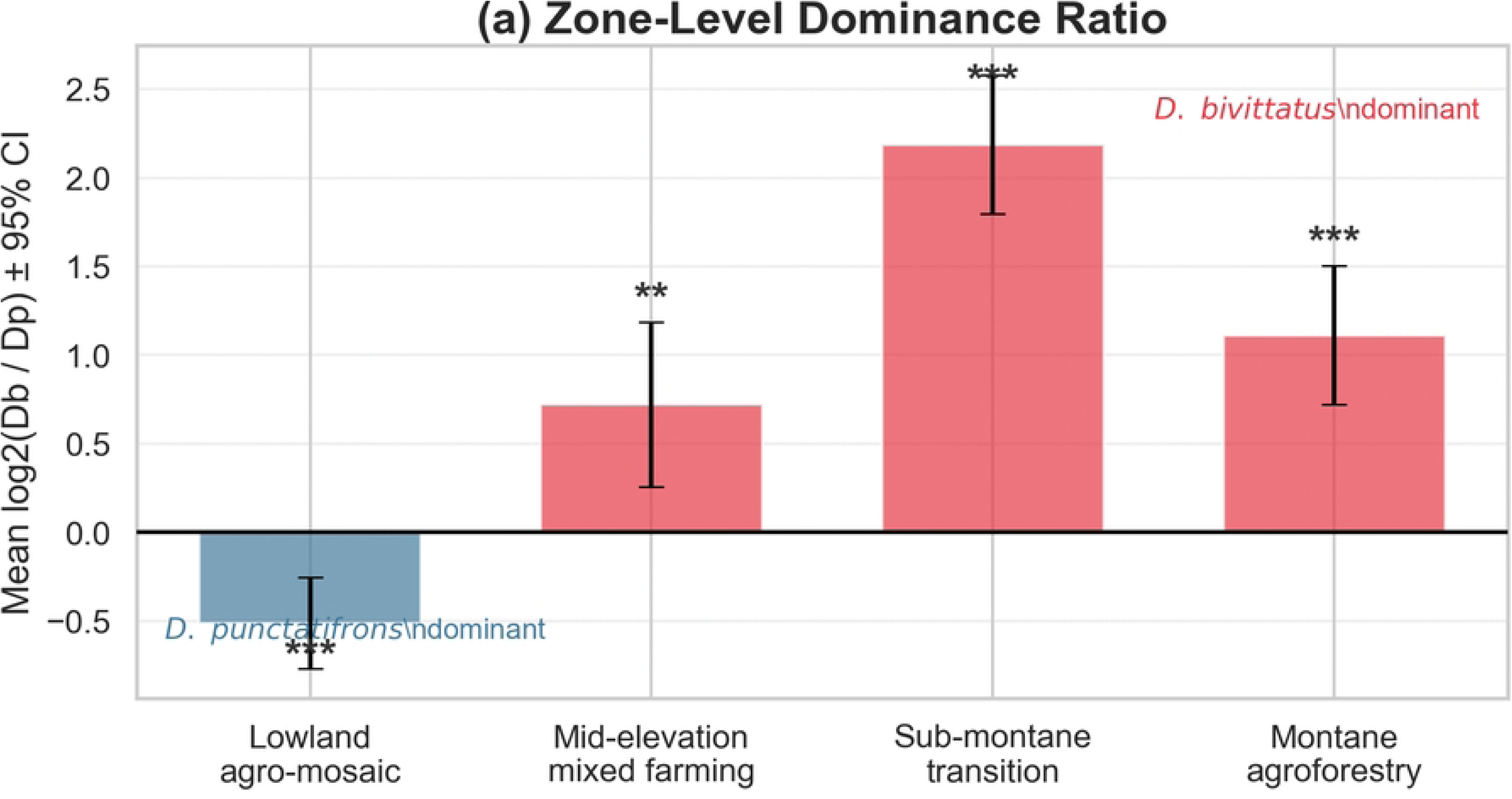
Dominance patterns between *D. bivittatus* and *D. punctatifrons* along the altitudinal gradient. Mean log_2_ dominance ratios across sites show the transition in species dominance from lowland to highland environments.

#### Niche overlap

Niche overlap was moderate along the altitudinal gradient. In contrast, seasonal overlap was high when pooled across sites, indicating that both species were active during similar months of the year (Fig. 9a–d). However, seasonal overlap declined with elevation. In the lowland zone, overlap remained high, but decreased through the transition and sub-montane and montane sites. This pattern indicates increasing ecological separation between the species under cooler highland conditions.

**Fig. 9.**
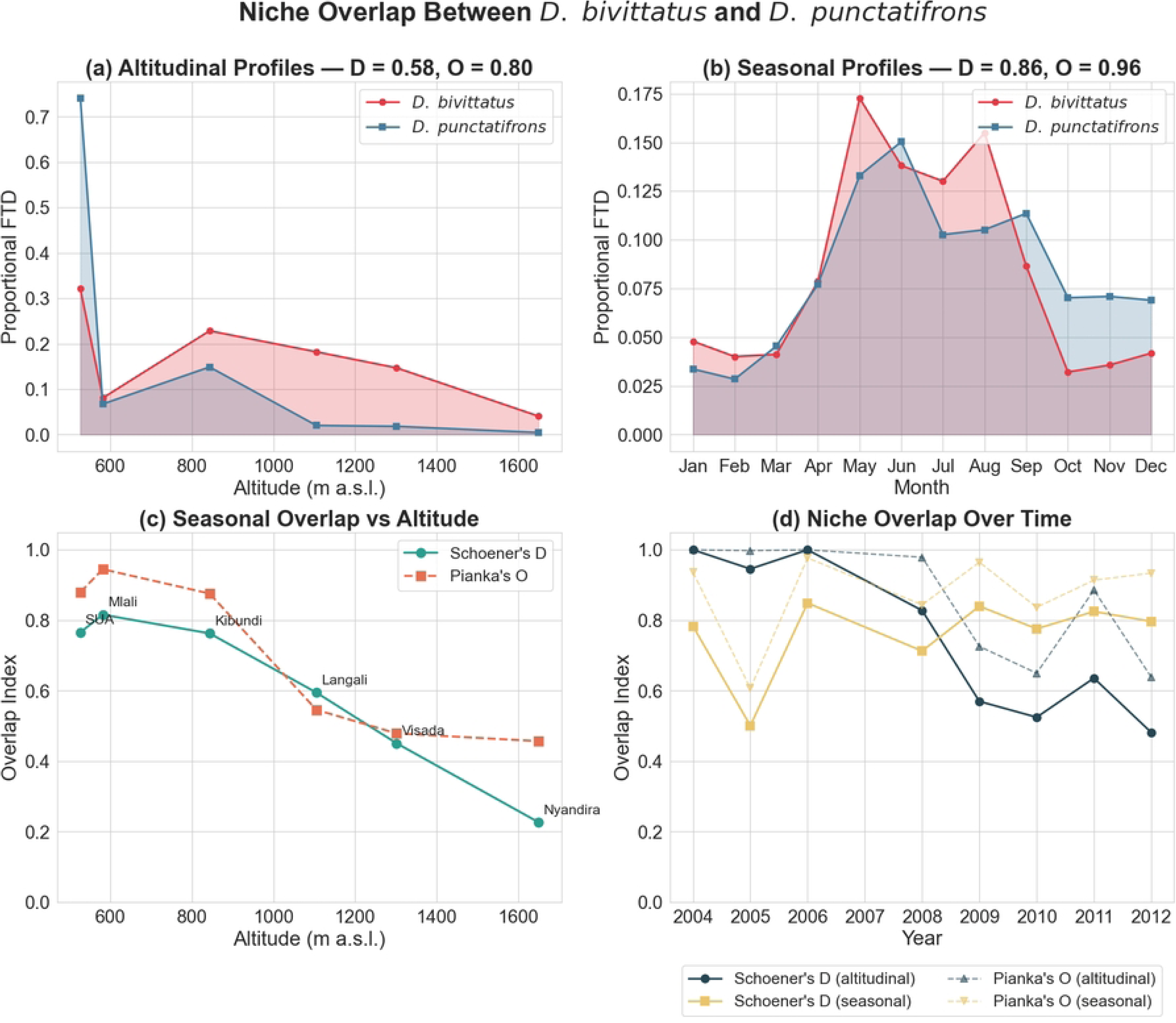
Niche overlap between *D. bivittatus* and *D. punctatifrons* across environmental gradients. Profiles of relative abundance across altitude and season show decreasing overlap at higher elevations.

## Discussion

This study demonstrates that environmental gradients along the Uluguru Mountains structure the ecological relationship between *D. bivittatus* and *D. punctatifrons*. Both species exhibited pronounced seasonal and interannual variation but differed in seasonal expression, spatial distribution, and responses to environmental conditions. These patterns indicate species-specific responses to altitude, climate, and agroecological context, highlighting the role of environmental gradients in shaping their population dynamics.

Despite their economic importance (1), the seasonal dynamics of *D. bivittatus* and *D. punctatifrons* remain insufficiently understood, with existing knowledge largely limited and fragmented (14,16,24), including studies from Morogoro that did not explicitly determine temporal variation (17,21,28). Long-term seasonal dynamics and comparative population ecology of the two species remain poorly resolved. Studies in Burkina Faso showed that *D. punctatifrons* can maintain distinct seasonal peaks even outside intensive vegetable systems, supported by wild hosts and non-crop habitats (16), indicating that its population dynamics extend beyond crop cycles and are shaped by broader landscape-level host availability. In Benin, populations of *D. punctatifrons* increased sharply from August, coinciding with changes in host availability and agroecosystem conditions (24). In contrast, the seasonal ecology of *D. bivittatus* remains even less well documented and is largely inferred from general surveys in cucurbit systems (for example, 14,15,17). Available evidence suggests that the activity of *D. bivittatus* may be influenced by seasonal temperature and host phenology (18), but these insights are derived from short-term observations and do not capture longer-term variability across environmental gradients. Clear seasonal peaks have also been reported in another cucurbit infester, *Z. cucurbitae*, which reaches maximum abundance during the dry season (July–September) (9).

Spatial patterns along the altitudinal gradient further structured species distribution. The abundance of the two species declined with increasing elevation, but more steeply in *D. punctatifrons*, resulting in a shift in dominance along the gradient. *D. punctatifrons* dominated in the warm lowlands, whereas *D. bivittatus* became increasingly dominant at higher elevations. This pattern suggests that altitude acts as an ecological filter, likely through its influence on temperature and habitat conditions. Previous reports on altitudinal gradients in Tanzania showed that the abundance of *D. bivittatus* declined significantly with increasing elevation (28). Similar elevation-driven structuring of fruit fly populations has been reported across multiple systems, where species distribution and community composition change along environmental gradients, for example, *Rhagoletis cerasi* L. in Turkey (38), tephritid assemblages in Papua New Guinea (39), fruit fly communities in Peru (40,41) and Mexico (42), as well as *Ceratitis capitata* (Wiedemann) in Greece (43).

Weather responses revealed further differences in climatic sensitivity between the two species. *D. punctatifrons* showed clear associations with temperature and rainfall, whereas *D. bivittatus* responded weakly, suggesting a lower dependence on short-term weather variability. This contrast indicates differences in ecological tolerance and environmental responsiveness. These findings are consistent with broader fruit fly studies, for example, *B. dorsalis* in Ghana, Kenya, and southern Africa (44–46), *Z. cucurbitae* in India (47), tephritid assemblages in Bangladesh (48), and *Bactrocera zonata* (Saunders) in India (49). We further found that niche relationships between the two species varied along the gradient. Seasonal overlap was high at low elevations, where both species co-occurred within similar temporal windows, but declined with increasing elevation, indicating greater separation under cooler conditions. This observation may be partly explained by the broader ecological distribution of *D. bivittatus*, which extends into relatively cooler regions of Africa (12). Together these findings suggest partial niche overlap combined with environmental differentiation, consistent with niche partitioning mechanisms observed in other fruit fly communities (50–53).

From our findings, landscape context and agroecological structure provide an additional layer of explanation for the observed patterns. The persistence of *D. punctatifrons* across seasons, including outside intensive cropping systems, shows the importance of alternative hosts and heterogeneous landscapes in sustaining populations. Along the Uluguru transect, agroecological zones integrate climatic conditions, host availability, and habitat structure, with warm, fruit-rich lowlands transitioning to cooler, less host-rich highland systems. The role of landscape configuration in shaping fruit fly populations has been demonstrated in other systems where habitat structure influences abundance, movement, and infestation patterns (54–59).

Overall, the ecology of *D. bivittatus* and *D. punctatifrons* is structured by interacting effects of altitude, climate, and landscape context across agroecological gradients. These findings provide new evidence that the two species respond differently to these drivers, resulting in contrasting distribution patterns between lowland and highland environments. This long-term comparative assessment further demonstrates that altitude acts as a key ecological filter, with host availability and agroecological conditions suggesting landscape-scale control of fruit fly population dynamics.

We recommend that control strategies should be adapted to agroecological zones, with greater emphasis on lowland systems where *D. punctatifrons* dominates and shows strong climatic responsiveness. Area-wide management approaches that incorporate wild hosts and non-crop habitats should be adopted. Finally, the observed variability across years and along environmental gradients underscores the need for long-term monitoring to improve prediction and guide sustainable pest management strategies.

## Conclusion

This long-term study demonstrates that altitude acts as the primary axis of ecological divergence between *D. bivittatus* and *D. punctatifrons* along the Uluguru Mountains transect. The two species differ fundamentally in seasonal strategy, climatic sensitivity, and altitudinal distribution, with agroecological zone emerging as the dominant predictor of abundance for both. The species dominance crossover at approximately 569 m a.s.l. and the progressive decline in seasonal niche overlap from 0.82 in the lowlands to 0.23 at 1,650 m confirm increasing ecological separation under cooler highland conditions. The stronger temperature dependence of *D. punctatifrons* suggests greater vulnerability to shifting thermal regimes under climate change. These findings support the adoption of zone-specific monitoring and management strategies, with priority given to lowland systems where both species co-occur at highest densities and where landscape-level integration of cultivated and wild cucurbit hosts into area-wide management will be most critical.

## Author’s contributions

Maulid W. Mwatawala (MWM): Conceptualization, Methodology, Investigation, Data Collection, Writing-original draft.

Marc De Meyer (MDM): Methodology, Resources, Data Collection, Writing-review & editing. Jackline Bakengesa (JB) and Mwajuma Zinga (MZ): Writing-review & editing.

Joseph O Ruboha (JOR): Formal analysis, Writing-review & editing

## Ethics

The research was conducted from 2004 and 2012, before the establishment of formal ethical review boards. The study involved only insect sampling and environmental observations and did not involve human participants or vertebrate animals. All procedures complied with institutional guidelines and accepted scientific practices applicable at the time of the study.

## Acknowledgements

The study was carried out in several phases between 2004 and 2012 and was supported by multiple funding sources, including the Belgian Development Cooperation through the Framework Programme with the Royal Museum for Central Africa (Projects F13 and S1_TNZ_IPM), the joint Food and Agriculture Organization / International Atomic Energy Agency (FAO/IAEA) Programme on Nuclear Techniques in Food and Agriculture through the Coordinated Research Project *“Resolution of Cryptic Species Complexes of Tephritid Pests to Overcome Constraints to the Sterile Insect Technique (SIT) Application and International Trade”*, the International Foundation of Science (Project C3956), and the Belgian Science Policy (Project MO/37/017).

## Declaration

Artificial intelligence (AI) tools were used solely for grammar checking, spelling correction, and improving language clarity. All analyses, interpretations, and scientific conclusions were conducted entirely by the authors.

